# BioLACE: unifying spatial geometry and marker priors for cohesive cell-type clustering in spatial transcriptomics

**DOI:** 10.64898/2025.12.01.691603

**Authors:** Haoran Qin, Yunfei Hu, Yuling Zhu, Jihoon Baek, Weiman Yuan, Shan Meltzer, Xin Maizie Zhou

**Affiliations:** Department of Computer Science, Vanderbilt University, Nashville, USA; Department of Biomedical Engineering, Vanderbilt University, Nashville, USA; Department of Pharmacology, Vanderbilt University, Nashville, USA

**Keywords:** Spatial Transcriptomics, Gene Expression, Cell-Type Clustering, Representation Learning, Marker Gene Priors

## Abstract

Spatial transcriptomics (ST) provides high-dimensional gene expression profiles together with spatial coordinates, enabling the reconstruction of tissue architecture at cellular resolution. While recent graph-based deep learning methods have advanced spatial clustering in ST, many rely on complex architectures that obscure interpretability and rarely integrate biological priors such as marker gene information. We introduce BioLACE, a scalable framework that unifies spatial structure, transcriptomic variation, and curated marker gene profiles within a shared Variational Autoencoder (VAE) latent space. BioLACEjointly optimizes three complementary objectives: (1) a VAE reconstruction loss to preserve transcriptional structure, (2) a graph Laplacian regularizer to enforce spatial smoothness, and (3) a temperature-scaled InfoNCE contrastive loss guided by marker-informed similarities. Applied to MERFISH hypothalamus, mouse spinal cord, and Slide-seq mouse cerebellum datasets, BioLACE achieves superior cell type clustering accuracy, well-defined biologically consistent boundaries, and interpretable latent representations, highlighting its generality and scalability for modern ST analysis. The source code and tutorials for BioLACE are publicly available at https://github.com/maiziezhoulab/BioLACE.

## 1 Introduction

Spatial transcriptomics (ST) [1] technologies profile gene expression while preserving each cell’s physical context, enabling analysis of how microenvironments shape cellular state. By jointly capturing molecular and spatial information, ST offers a unique view into tissue organization, cell–cell communication, and developmental patterning. A central computational challenge is to delineate coherent cell populations and tissue domains that respect both transcriptional similarity and spatial topology. Although recent methods have improved spatial clustering accuracy and scalability [2], many still struggle to achieve three critical goals simultaneously: geometric fidelity to tissue structure, discriminative clustering of biologically distinct types, and interpretability grounded in known molecular programs.

### Progress and remaining challenges

Early statistical frameworks emphasized spatial smoothness and neighborhood consistency. BayesSpace [3] introduced a Bayesian model that refines subspot resolution by encouraging adjacent spots to share labels, while BANKSY [4] augmented each spot’s features with neighborhood averages and gradients to stabilize domain boundaries. These methods capture local spatial continuity but rely on predefined priors rather than learned representations. Deep learning models extend this idea by jointly embedding molecular and spatial signals into a unified latent space. SpaGCN [5] integrates gene expression, spatial coordinates, and histology through graph convolution to detect spatially variable genes, and SEDR [6] couples an autoencoder with a variational graph model to encode latent spatial relationships. ADEPT [7] enables better domain identification by using region markers with iterative GAT-based clustering. More recent frameworks, including GraphST [8] incorporate self-supervision or contrastive objectives to improve boundary definition and scalability. Yet broad benchmarking studies reveal persistent trade-offs: Bayesian models are transparent but limited in scale and flexibility, whereas deep graph networks achieve higher accuracy at the cost of interpretability and explicit biological supervision. In particular, these representations are often *agnostic* to prior biological knowledge—treating all genes as equally informative despite decades of curated marker-gene literature.

### Motivation for biological priors in cell-type clustering

Marker genes encode reproducible transcriptional programs that define molecularly distinct cell types across tissues and modalities; importantly, these signatures capture cell-intrinsic identity (as exploited by cell-type classifiers such as CellAssign [9] and Garnett [10]) rather than higher-order tissue domains, which often contain mixtures of multiple cell types shaped by local microenvironmental gradients. Many existing spatial methods incorporate marker information only for *post hoc* annotation, [9, 11], leaving the representation itself driven purely by expression and geometry. In contrast, we treat marker programs as soft relational constraints during training, encouraging cells that co-express shared identity signatures to occupy nearby positions in latent space while still allowing unannotated subpopulations to emerge. By guiding representation learning at the level of cell types—rather than directly imposing domain structure—BioLACE leverages decades of curated biological knowledge to produce embeddings that are both interpretable and compatible with heterogeneous spatial patterns.

### Contrastive learning for spatial omics

Contrastive learning optimizes pairwise relationships by pulling similar samples together and pushing dissimilar ones apart in latent space [12, 13]. In spatial omics, such objectives capture cross-context similarity—for example, linking spatially distant cells that express the same molecular program, or separating neighboring spots belonging to distinct microenvironments. BioLACE adapts this paradigm by defining positive and negative pairs from marker-informed and spatial cues, transforming prior knowledge into a differentiable relational signal that refines the embedding rather than merely labeling it after training.

### Balancing spatial coherence and biological discrimination

Spatial regularization enforces local continuity, ensuring nearby cells share similar latent representations, but excessive smoothness can blur boundaries between distinct cell types. Conversely, contrastive learning sharpens biological distinctions by separating dissimilar profiles, yet overly strong contrastive pressure can fragment continuous spatial gradients. Our ablation on MERFISH hypothalamus data (Sec. 3.3) demonstrates that co-training these objectives, as well as carefully balancing their relative strength, achieves the best of both: anatomically coherent yet biologically distinct embeddings. BioLACE thus resolves the classic over-smoothing [14] versus over-segmentation dilemma through a temperature-controlled joint optimization of spatial and biological constraints.

### Our contribution

We introduce BioLACE, a biologically guided deep clustering framework for spatial transcriptomics. BioLACE integrates three complementary objectives within a shared variational latent space: (i) a variational autoencoder (VAE) reconstruction loss to preserve transcriptional structure, (ii) a graph Laplacian regularizer enforcing spatial coherence, and (iii) a marker-aware contrastive loss encoding biological priors as relational constraints. By embedding biological knowledge directly into training, BioLACE produces latent representations that are spatially smooth, biologically interpretable, and robust across ST platforms. Applied to Slide-seq V1/V2 cerebellum [15, 16] and MERFISH spinal cord datasets, BioLACE reconstructs known architecture, enhances marker co-expression, and reveals fine-grained domains missed by reference-based annotation.

## 2. Methods

### Design goal

BioLACE learns a latent space where (i) transcriptionally similar cells are close, (ii) spatial neighbors remain coherent, and (iii) curated biological programs define interpretable axes of separation. In practice, this means encoding three complementary signals—expression, geometry, and biology—during representation learning, not as post hoc annotation.

### 2.1 Data model and notation

#### Spatial transcriptomics as structured data

A spatial transcriptomics (ST) dataset consists of thousands of measurement points (spots or cells), each defined by a gene expression profile and a spatial coordinate marking its position within the tissue. Our goal is to transform this high-dimensional, spatially structured data into a compact latent representation that preserves both biological similarity and anatomical proximity.

#### Notation

Let **X** ∈ ℝ^*N ×G*^ denote the log-normalized gene expression matrix, where each of *N* cells is profiled across *G* genes. The spatial coordinates **s**_*i*_ = (*x*_*i*_, *y*_*i*_) for each cell form **S** ∈ ℝ^*N ×*2^, which defines a geometric map of the tissue. From these coordinates we construct a *k*-nearest-neighbor graph 𝒢 = (𝒱, ℰ ), with edge weights

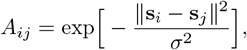

so that nearby cells have strong connections and distant ones have weak or zero weights. The adjacency matrix **A** encodes local tissue topology; the degree matrix **D** = diag( ∑_*j*_ *A*_*ij*_) gives the *graph Laplacian* **L** = **D A**, which penalizes large latent differences between neighboring cells—acting as a spatial smoothness operator.

#### Latent representation

Each cell *i* is mapped to a *d*-dimensional embedding **z**_*i*_ ∈ ℝ^*d*^, forming **Z** ∈ ℝ^*N ×d*^. Distances in this space reflect biological and spatial relatedness. Clusters are obtained by applying Leiden clustering [17] to **Z**, producing cluster labels *c*_*i*_ ∈*{*1, …, *K}*.

#### Marker genes

When prior biological knowledge is available—such as expectations about which cell types should appear in a given tissue or marker panels curated from external atlases (e.g., the Allen Brain Atlas [18])—we construct the marker set ℳ ⊆ {1, …, *G*} by filtering genes according to these priors. Such curated markers act as soft biological anchors, promoting proximity among cells that co-express shared transcriptional programs while still permitting the discovery of unannotated states. When no reliable priors exist, ℳis estimated internally: pseudo-clusters are generated via Leiden [17] clustering on raw input gene expression, and cluster-enriched genes are identified using Wilcoxon rank-sum testing. The union of top-ranked genes across clusters defines a surrogate marker set ℳ_pseudo_. Substituting ℳ_pseudo_ with experimentally validated markers yields improved biological purity and interpretability while maintaining comparable geometric coherence (see Sec. 3.2).

### 2.2 From data to objectives: modeling assumptions

BioLACE is built on three complementary assumptions: (1) gene expression lies on a smooth manifold that can be denoised and compressed via a generative latent model. (2) nearby spatial locations share microenvironmental context, so embeddings should maintain neighbor relations unless contradicted by expression. (3) curated marker sets encode reproducible transcriptional programs that can act as soft relational constraints guiding the embedding. These principles motivate three corresponding losses—a VAE reconstruction term, a Laplacian spatial regularizer, and a marker-aware contrastive objective—optimized jointly.

### 2.3 Backbone: Variational autoencoder for transcriptomic structure

A variational autoencoder [19, 20] (VAE) captures the global expression manifold while denoising technical noise. It provides a nonlinear, probabilistic embedding that preserves gene–gene correlations and enables downstream integration of spatial and biological regularization. For each cell *i*:

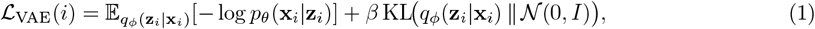

where *β* controls the information bottleneck. This step anchors BioLACE in a likelihood-based foundation before adding spatial and biological constraints.

### 2.4 Spatial graph alignment for geometric coherence

To ensure spatial coherence, we penalize latent separation of neighboring cells using the Laplacian regularizer:

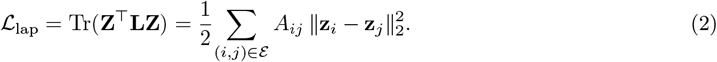

This encourages smooth transitions within tissue neighborhoods while allowing sharp boundaries when supported by expression. The Laplacian is differentiable, convex in **Z**, and compatible with stochastic optimization—making it a natural choice over discrete graph-cut or total variation penalties.

### 2.5 Encoding biological priors through marker-guided similarity

Curated marker genes ℳ are treated as soft relational constraints rather than fixed labels. They are translated into cell-cell pairwise similarities that bias the embedding toward biologically consistent neighborhoods.

#### Spatially smoothed features

We stabilize sparse expression signals by aggregating *H*-hop spatial neighbors:

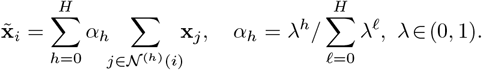

This smoothing mitigates dropout noise while preserving local context. *λ* controls geometric decay of neighbor influence: small values emphasize local structure (sharp boundaries), while larger values propagate information across multiple hops to denoise sparse signals.

#### Two complementary similarities

We compute two cosine-based similarities for each cell pair (*i, j*):

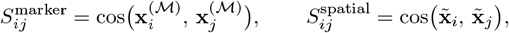

where *S*^marker^ measures co-expression of curated marker programs and *S*^spatial^ captures smoothed neighborhood context.

#### Global fusion and pair binning

To emphasize pairs that are jointly marker-consistent and spatially aligned, we form a fused similarity:

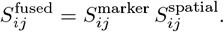

Applying two-centroid *k*-means to 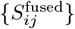 yields global thresholds *τ* ^+^ and *τ* ^−^. We then construct two global pair sets,

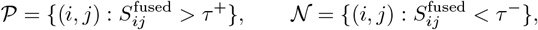

representing biologically supported “similar” and “dissimilar” pairs. All remaining pairs are ignored. These sets constitute the sole supervision for the contrastive component of BioLACE. The next section describes how they are incorporated into the loss.

### 2.6 Contrastive learning with mined positive and negative pairs

#### Formulation and departure from standard InfoNCE

Although 𝒫 and 𝒩 are defined globally, the contrastive loss evaluates them one cell at a time to enable per-cell normalization. For a given cell *i*, we collect all mined partners involving that cell:

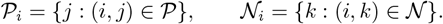

Thus, 𝒫_*i*_ and 𝒩_*i*_ are simply the subsets of globally mined pairs that relate to cell *i*, specifying which cells it should be pulled toward (marker-consistent, spatially aligned) and pushed away from (reliable counter-examples). This differs from classical InfoNCE [12], which treats *all* other samples in a batch as negatives. By restricting comparisons to mined pairs, BioLACE avoids penalizing uncertain relationships and transforms curated markers into a precise, semi-supervised relational signal, analogous to supervised contrastive learning [21].

#### Contrastive objective

Let *s*_*ij*_ = sim(**z**_*i*_, **z**_*j*_) denote cosine similarity in latent space. For each positive pair (*i, j*) ∈ 𝒫, we encourage *s*_*ij*_ to be large by contrasting it against the mined negatives 𝒩 _*i*_:

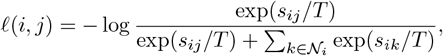

where *T* is a temperature that controls contrastive sharpness. We use a symmetric form that counts both directions (*i, j*) and (*j, i*):

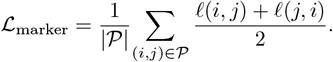

#### Interpretation

Only mined negatives appear in the denominator, focusing repulsion on reliable dissimilar examples and reducing gradient noise from ambiguous pairs. The temperature *T* ∈ [0.5, 2.0] (decayed during joint training; Sec. 2.7) controls the strength of discrimination, ensuring stable integration with the spatial Laplacian term.

### 2.7 Joint training objective and schedule

The full objective combines all terms:

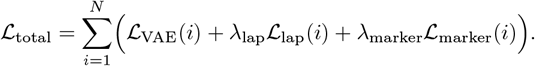

Training proceeds in two phases: (1) Warmup (5–20 epochs) with small (*λ*_lap_, *λ*_marker_) to let the VAE stabilize, and (2) Joint optimization, where (*λ*_lap_, *λ*_marker_) are linearly ramped to their target values while the temperature *T* decays following a cosine schedule. This schedule avoids premature geometric forcing and gradually harmonizes spatial and biological constraints.

### 2.8 Clustering and evaluation

Latent embeddings **Z** are clustered with Leiden (resolution tuned by Silhouette). Evaluation metrics include ARI and V-measure against ground truth (or pseudo-ground truth) annotations, AUROC/AUPRC of marker modules, within-cluster coexpression (Spearman *ρ*), Moran’s *I* [22] for spatial autocorrelation, and graph modularity.

### 2.9 Complexity and implementation

All objectives support stochastic mini-batching for efficient GPU training. Constructing the *k*-nearestneighbor spatial graph scales as *O*(*N* log *N* ) (or *O*(*kN* ) with prebuilt indices), and each training epoch costs

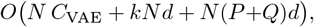

where *C*_VAE_ denotes the per-sample encoder–decoder cost, *d* the latent dimension, and *P, Q* the average number of positive and negative pairs used in the contrastive term. For fixed *k, d, P*, and *Q*, both runtime and memory scale linearly with the number of cells or spots *N* . The framework is implemented in PyTorch with Scanpy/Squidpy preprocessing, using neighborhood sampling to avoid *O*(*N* ^2^) pair storage.

## 3 Results

### 3.1 Slide-seq cerebellum: spatial fidelity and functional coherence

#### Dataset overview

We evaluated BioLACE on the mouse cerebellum Slide-seq datasets, a high-resolution spatial transcriptomic assay that captures transcript counts from thousands of 10 *µ*m beads across the tissue surface. The cerebellum is particularly well-suited for benchmarking because it features well-defined laminar organization and distinct molecular zones. Slide-seq V1 comprises a compact sagittal section containing clear Purkinje, granule, and myelinated white-matter layers, offering a stringent test of geometric fidelity. Slide-seq V2 spans a broader field of view with interleaved neuronal, glial, and vascular microenvironments, challenging methods to maintain biological coherence while preserving fine-grained spatial boundaries. Together, these datasets provide complementary benchmarks: V1 emphasizes structural precision, whereas V2 stresses functional interpretability across complex cellular niches.

#### Clustering accuracy and neighborhood agreement

Compared with BANKSY and SpiceMix, and using RCTD [23]-derived reference-based cell-type annotations as pseudo-ground truth, BioLACE achieves superior performance on both datasets. In V1, BioLACE attains an ARI of 0.445 and V-measure of 0.377 (BANKSY 0.188*/*0.160, SpiceMix 0.182*/*0.313); in V2, it reaches an ARI of 0.446 and V-measure of 0.501 (BANKSY 0.313*/*0.357, SpiceMix 0.152*/*0.349) (Fig. 2a). Representative cell types—including the Granular layer, Astrocytes, and Choroid Plexus—show strong clustering correspondence between RCTD and BioLACE (Fig. 2b). Neighbor-enrichment analysis using Squidpy further demonstrates consistently higher same-label probabilities among spatial neighbors for BioLACE, indicating that the Laplacian term anchors the latent manifold to tissue topology without oversmoothing boundaries (see Supplementary Fig. 1).

**Fig. 1.**
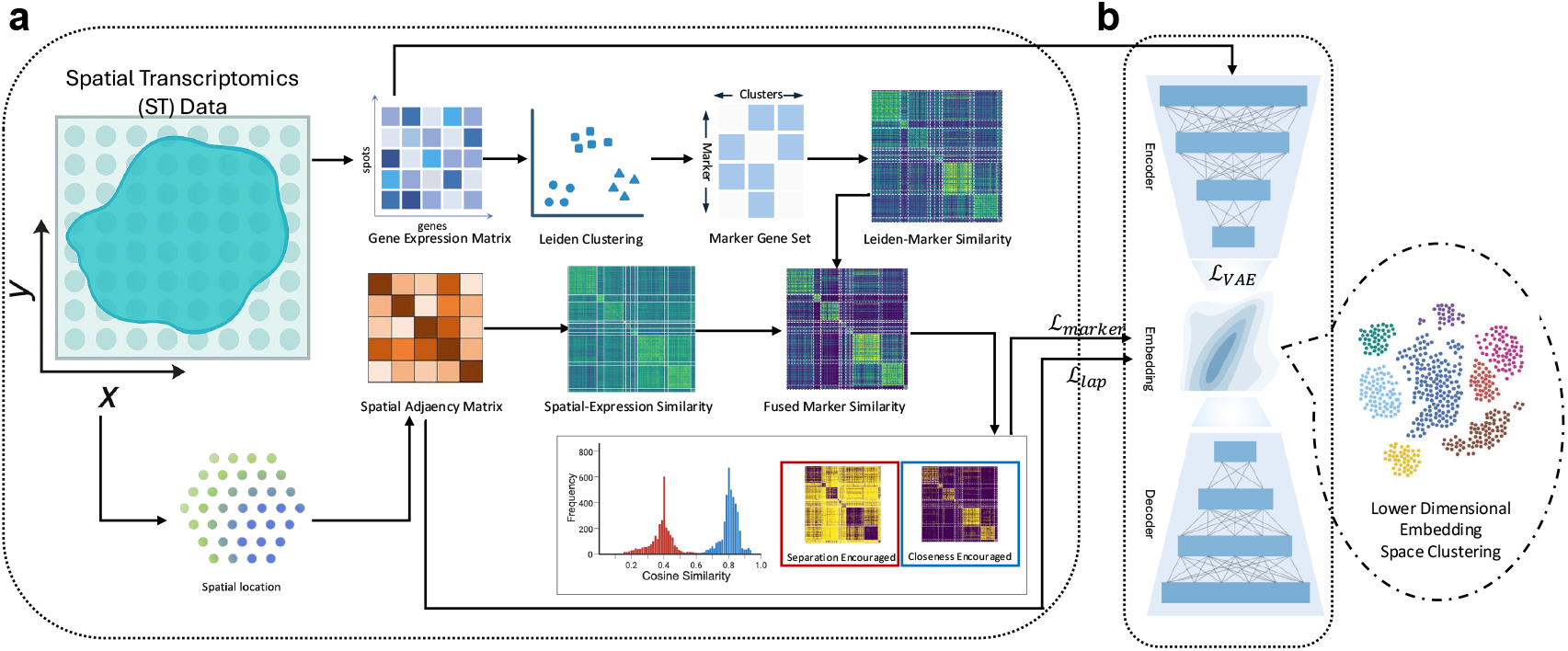
Overview of the BioLACE pipeline. **(a)** Spatial transcriptomics (ST) data provide geneexpression profiles for each spot together with its (*x, y*) coordinates. From these we construct a gene-expression matrix and a spatial *k*-nearest-neighbor graph. An initial Leiden clustering, followed by Wilcoxon testing, yields a pseudo–marker-gene set for each preliminary cluster. These markers define a marker-based similarity matrix, which is combined with a spatially smoothed expression–similarity matrix via a Hadamard product. The fused similarity exhibits distinct highand low-similarity modes; two-centroid *k*-means identifies the reliable positive and negative cell pairs used for contrastive learning, where the positive pairs are encouraged to be close, and negative pairs are encouraged to separate in the latent space. **(b)** BioLACE integrates the three processed inputs—gene expression, spatial adjacency, and marker-derived similarity—through a variational autoencoder (VAE) backbone with two regularizers. The VAE thus learns a low-dimensional embedding by co-optimizng three losses: a reconstruction loss ℒ_VAE_ preserving transcriptomic structure; a Laplacian loss ℒ_lap_ derived from the spatial adjacency matrix that maintains neighborhood continuity; and a marker-guided contrastive loss ℒ_marker_ enforcing the mined pairwise relations. Clustering the resulting embedding produces spatially coherent and biologically interpretable cell type clusters.

**Fig. 2.**
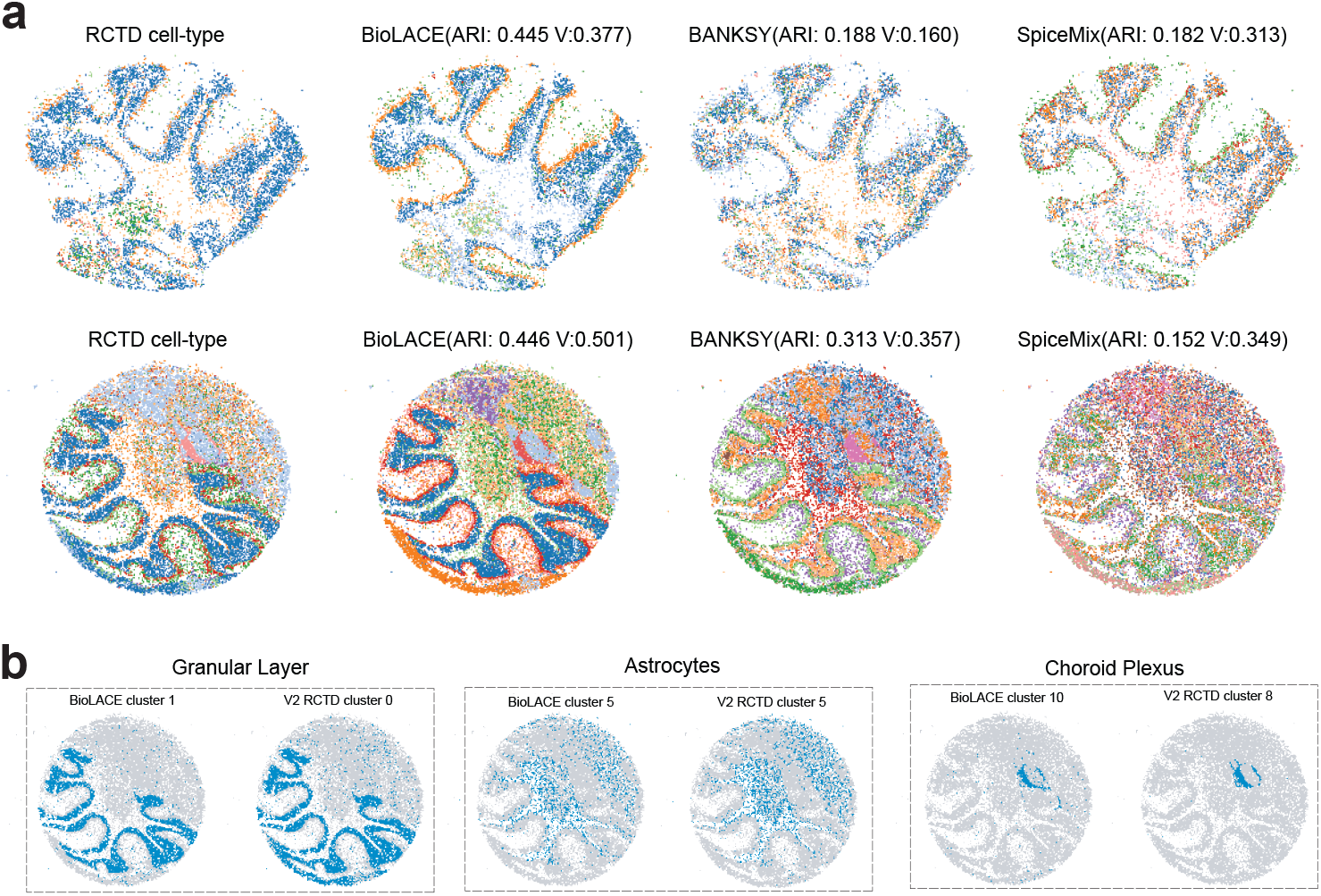
Clustering on Slide-seq V1/V2 and agreement with RCTD Cell-Type Labels. (**a)** V1 (top): spatial maps for RCTD, BioLACE, BANKSY, and SpiceMix; BioLACE shows strong alignment with RCTD labels and reproduces laminar boundaries and myelin tracts. V2 (bottom): spatial maps across methods; BioLACE shows strong alignment with RCTD labels, and separates neuronal, glial, and vascular territories while preserving fine Purkinje subdomains. **(b)** Per-cluster matching between BioLACE and RCTD for Granular Layer, Astrocytes, and Choroid Plexus regions in V2, illustrating one-to-few correspondences rather than many-to-many splits.

#### Spatial autocorrelation reveals anatomically faithful clustering

Moran’s *I* quantifies how consistently a gene’s expression co-varies among nearby spatial locations—high values indicate genes confined to specific tissue domains. Because BioLACE does not explicitly optimize this statistic, elevated Moran’s *I* serves as an independent test of biological realism: if our clusters reflect genuine anatomy, their defining genes should naturally exhibit strong spatial autocorrelation.

Across both Slide-seq datasets, this pattern holds strikingly. In the cerebellar V1 tissue, genes with the highest Moran’s *I*, including *Plp1* and Purkinje-associated *Car8, Calb1, Pcp2*, and *Pcp4* (Fig. 3a), align precisely with BioLACE clusters that trace the myelin (Oligodendrocyte) tract and Purkinje layer through visual analysis (Fig. 3b), and quantitative analysis of inside- and outside-cluster co-expression shows that BioLACE preserves functional co-expression of canonical pairs such as *Pcp2* –*Pcp4* (*ρ*=0.65) and *Plp1* –*Mbp* (*ρ*=0.20) (Fig. 3c), further showing how BioLACE recovers clusters with strong biological explainability [18].

**Fig. 3.**
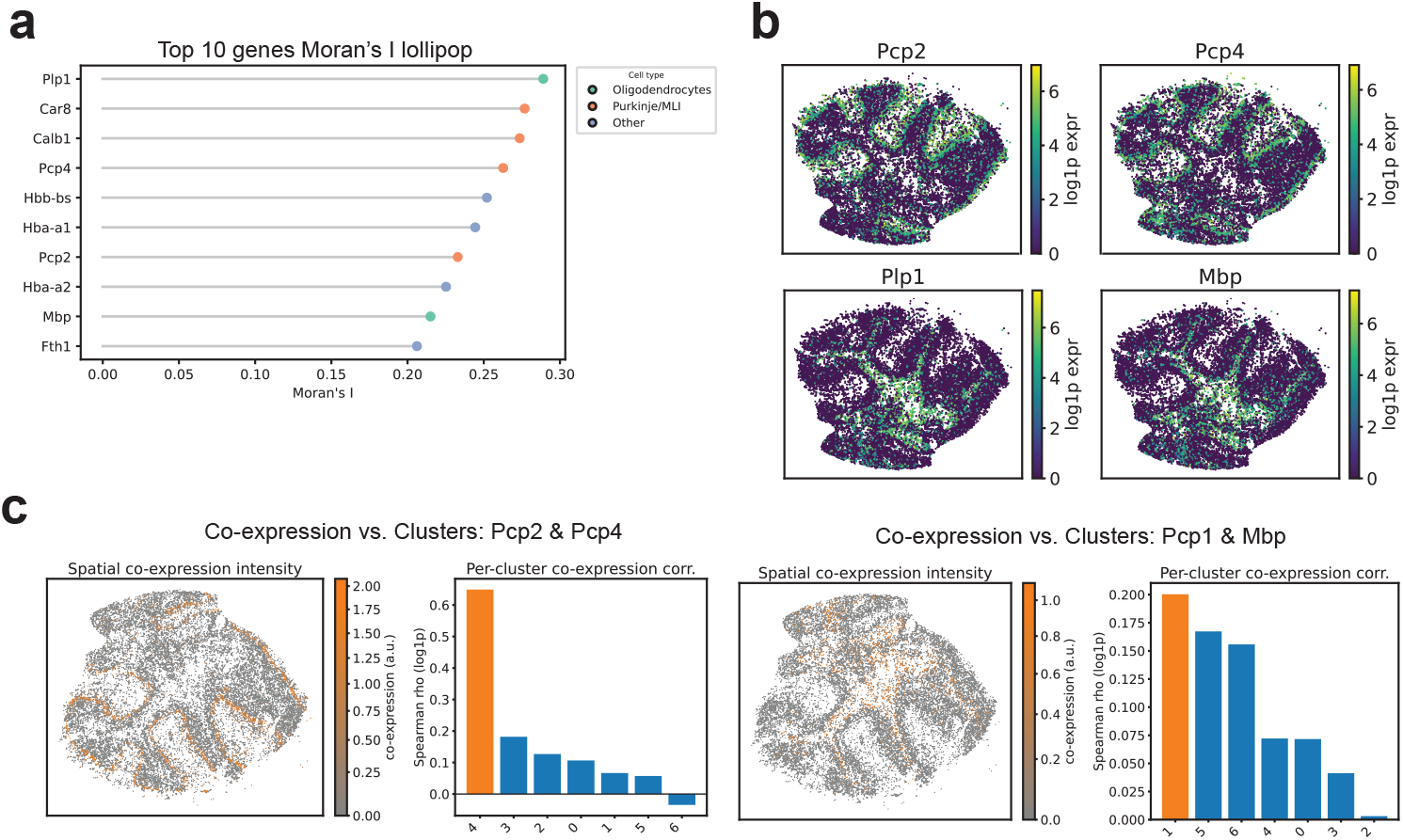
Slide-seq V1: Moran’s *I* and co-expression confirm laminar precision of BioLACE. **(a)** Lollipop plot ranking Moran’s *I* across all detected genes highlights *Plp1* (myelin, *I*=0.289), *Car8* and *Calb1* (Purkinje-associated), and *Pcp4* (Purkinje, *I*=0.263), followed by hemoglobin genes (*Hbb-bs, Hba-a1, Hba-a2* ) and myelin markers (*Mbp*). **(b)** Spatial expression overlays show that high–Moran’s *I* markers align precisely with BioLACE clusters: *Pcp2* /*Pcp4* localize to cluster 4 (Purkinje), while *Plp1* /*Mbp* localize to cluster 1 (oligodendrocytes). **(c)** Within-cluster co-expression (Spearman *ρ*) further validates functional coherence: *Pcp2* –*Pcp4* (*ρ*=0.649) in the Purkinje domain and *Plp1* –*Mbp* (*ρ*=0.200) in the myelin domain, both markedly higher than their out-of-cluster correlations. Together, these statistics show that BioLACE reconstructs anatomically faithful laminar structure without imposing spatial smoothing.

Additional quantitative analysis of Moran’s *I* marker in BioLACE’s clusters can be found in the Supplementary Table 1, detailing metrics such as AUROC.

The broader and more heterogeneous V2 dataset further reinforces this connection between spatial auto-correlation and biological structure through similar analysis; full Moran’s *I* and co-expression visualizations for V2 are provided in Supplementary Fig. 2.

#### BioLACE resolves a noradrenergic microdomain missed by label transfer

Beyond recapitulating known cell types, BioLACE uncovers a compact cluster co-expressing the full catecholamine biosynthetic and handling program—an interpretable population that was missed by RCTD’s reference-based label transfer. The cluster shows coordinated high expression of *Th, Ddc, Dbh, Slc18a2*, and *Slc6a2*, forming a complete noradrenergic program: *Th* and *Ddc* support catecholamine synthesis, *Dbh* converts dopamine to norepinephrine, *Slc18a2* packages monoamines into vesicles, and *Slc6a2* mediates norepinephrine reuptake (Fig. 4a-b). Mean expression (log1p) inside vs. outside the cluster is markedly elevated for *Slc18a2* (2.12 vs. 0.04), *Slc6a2* (2.08 vs. 0.04), *Dbh* (1.81 vs. 0.03), *Th* (1.27 vs. 0.03), and *Ddc* (1.35 vs. 0.04) (Fig. 4c), in contrast, the dopaminergic transporter *Slc6a3* is near baseline, ruling out a dopaminergic identity. This combination defines a bona fide noradrenergic microdomain that RCTD’s label transfer fails to resolve, underscoring BioLACE’s ability to resolve fine-grained states that are collapsed in reference-based mappings (Fig. 2c).

**Fig. 4.**
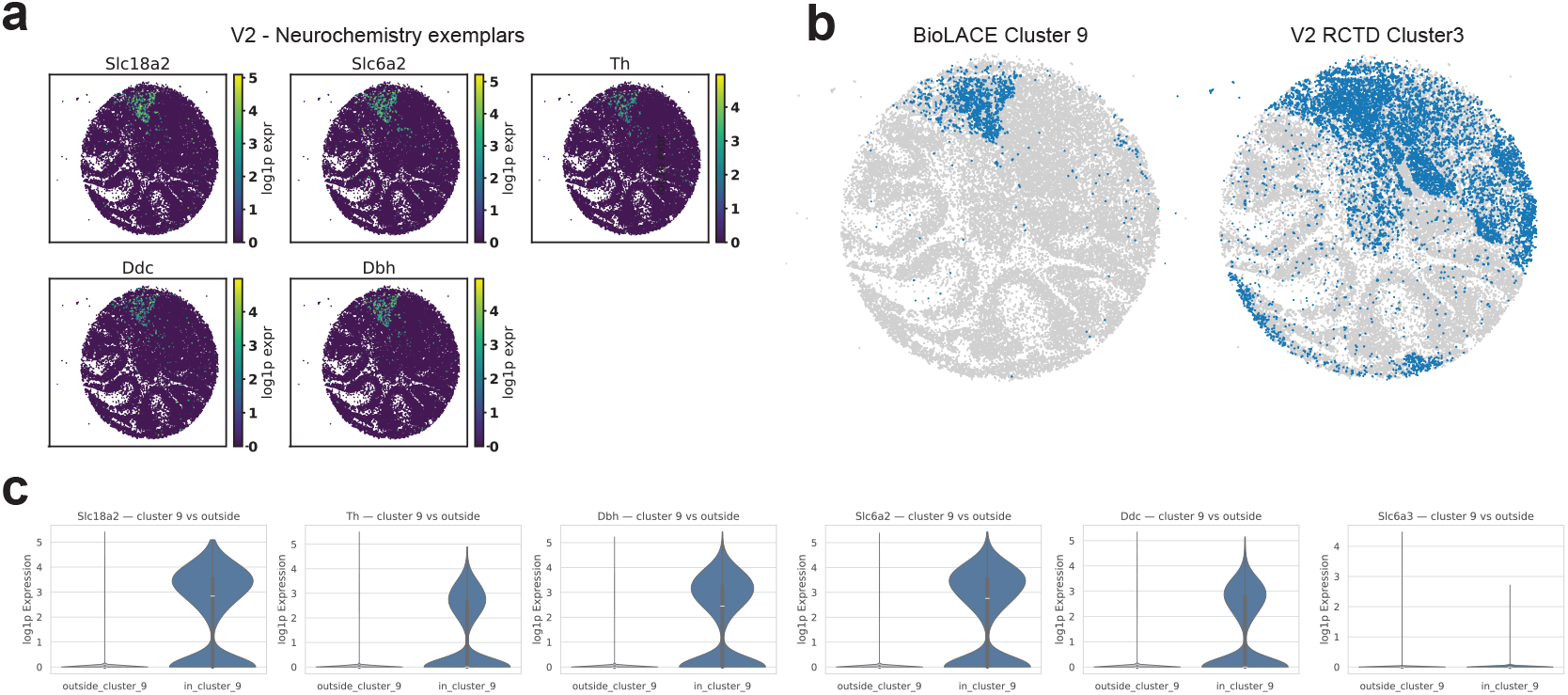
BioLACE reveals a noradrenergic microdomain not recovered by label transfer. **(a)** Spatial overlays reveal a compact cluster with coordinated expression of the canonical noradrenergic program—*Th, Ddc, Dbh, Slc18a2*, and *Slc6a2* —indicating a discrete catecholaminergic domain. **(b)** Comparison of BioLACE cluster 9 with RCTD’s corresponding label shows that BioLACE isolates this microdomain cleanly, whereas RCTD merges it into a broader neighboring region. **(c)** Inside–outside expression contrasts (log1p) confirm strong enrichment for all five genes (e.g., *Slc18a2* 2.12 vs. 0.04; *Slc6a2* 2.08 vs. 0.04; *Dbh* 1.81 vs. 0.03; *Th* 1.27 vs. 0.03; *Ddc* 1.35 vs. 0.04), while dopaminergic *Slc6a3* remains essentially absent, supporting a noradrenergic identity. Together, these results show that BioLACE resolves fine-grained, biologically coherent microstructures that are not captured by label-transfer–based annotation.

#### Cross dataset synthesis

Across Slide-seq V1/V2, BioLACE consistently achieves both geometric precision and biologically grounded cluster recovery. In V1, BioLACE sharply delineates laminar and myelinated domains, outperforming state-of-the-art spatial clustering methods in ARI, V-measure, and spatial neighborhood agreement—reflecting its explicit regularization for spatial contiguity. In V2, where neuronal, glial, and vascular microenvironments interleave at fine scales, BioLACE further demonstrates high marker-based interpretability and functional coherence. Notably, BioLACE detects biologically coherent microdomains—including the catecholaminergic niche—missed by RCTD label transfer, highlighting its ability to recover fine-grained structure that aligns with canonical marker programs.

### 3.2 Biological priors on MERFISH spinal cord data yield functionally interpretable embeddings

We next evaluated BioLACE on a high-resolution MERFISH dataset of the mouse spinal cord [24], which profiles both neuronal and non-neuronal populations spanning dorsal, ventral, and vascular compartments. The spinal cord provides an ideal system to test whether incorporating marker-gene priors into representation learning enhances not only clustering performance but also downstream functional interpretability in spatially complex tissue.

#### Integrating curated biological priors

We curated a panel of 43 canonical marker genes from spinal cord atlases and prior single-cell studies (see Supplementary Table 2 for the full list and rationale). These markers collectively capture excitatory and inhibitory neurons, glial and vascular populations, and ependymal support cells—e.g., *Slc17a8, Tac2, Nmu*, and *Prkcg* for dorsal horn excitatory subtypes; *Chat* and *Slc18a3* for ventral motoneurons; *Pdgfra* and *Opalin* for oligodendrocyte progenitors; and *Abcc9, Flt1*, and *Eng* for vascular and pericytic niches. Embedding these curated priors directly into the contrastive objective allows their co-expression structure to guide latent geometry, coupling biological interpretability with spatial representation learning.

#### Leveraging biological priors substantially improves BioLACE accuracy

Across the full spinal cord tissue, BioLACE achieved an ARI score of 0.271, outperforming BANKSY (0.229) and SpiceMix (0.117). When the curated biological prior was activated, modularity further increased to 0.437 (Fig. 5a), demonstrating that prior-informed contrastive learning enhances the alignment between clusters and true cell types. Spatial maps reveal clearer boundaries among major populations—including dorsal horn excitatory and inhibitory neurons, motoneurons, oligodendrocytes, astrocytes, and vascular/meningeal cells—recapitulating known laminar and perivascular organization (Fig. 5a). This improvement also manifests in labeling accuracy: using pseudocluster-derived markers, accuracy and recall increased from 73.36% and 56.54% (without priors) to 87.62% and 85.65% (with priors). Fig. 5b shows that incorporating priors substantially sharpens within–cell-type similarity in the model. These results highlight BioLACE’s strength when reliable biological priors are available. Supplying expected cell types and their marker programs guides the model to incorporate known tissue structure, producing more faithful and interpretable cell-type partitions.

**Fig. 5.**
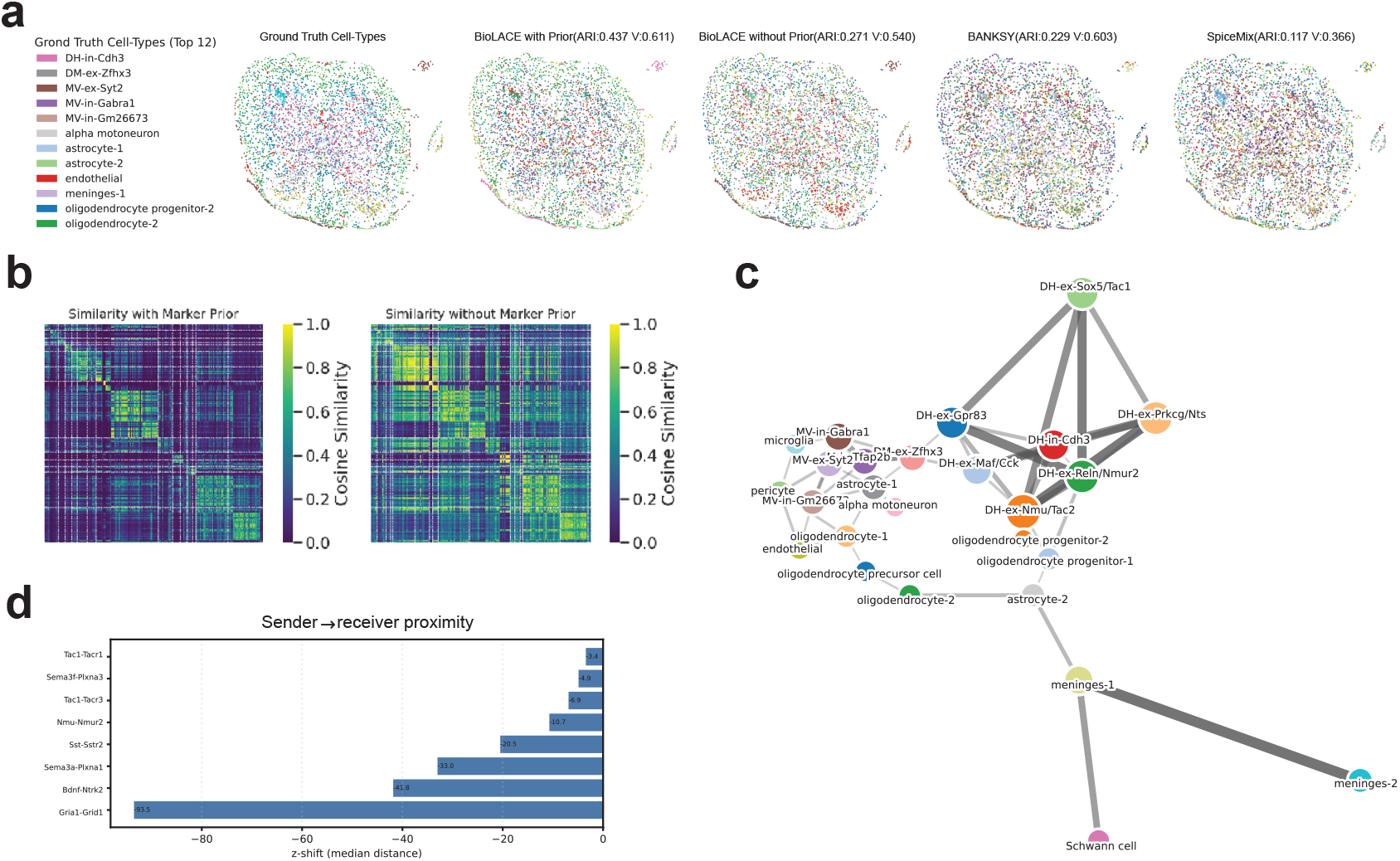
Biological priors enhance spatial and functional coherence on MERFISH spinal cord data. **(a)** Spatial cluster maps for BANKSY, SpiceMix, and BioLACE with and without marker priors (ARI = 0.437 vs. 0.270), filtered to the 12 largest ground truth cell types for clearer visualization. **(b)** Marker similarity matrices comparing prior-guided and unguided embeddings show substantially improved accuracy (87.6%) and recall (85.7%) with priors. **(c)** Spatial proximity graph highlights clear separation of dorsal horn (excitatory/inhibitory), ventral motoneuron, glial, and vascular compartments, recapitulating known laminar architecture. **(d)** Ligand–receptor network analysis shows spatially compact communication modules, with canonical signaling axes (*Bdnf–Ntrk2, Sema3a–Plxna1, Sst–Sstr2* ) confined to biologically appropriate regions. Together, these results demonstrate that integrating curated marker priors into contrastive learning yields embeddings with improved spatial fidelity and biologically meaningful communication structure.

#### Preserving anatomical context through proximity networks

To assess whether BioLACE preserves true tissue architecture rather than producing spatially smoothed artifacts, we analyzed neighborhood graphs constructed from BioLACE’s latent representation with Squidpy [25] (Fig. 5c). The resulting proximity networks revealed co-localization patterns that align with well-established spinal organization, including the dorsal–ventral laminar segregation of neuronal subtypes and their associated microcircuits, as reported in large-scale molecular atlases of the adult mouse spinal cord [26].

Excitatory subnetworks in the dorsal horn appeared with the expected layered structure: *DH-ex-Gpr83* consistently neighbored *DH-ex-Reln/Nmur2*, while *DH-ex-Nmu/Tac2* co-occurred with *DH-ex-Prkcg/Nts*, reflecting known functional pairings. Inhibitory populations such as *DH-in-Cdh3* showed balanced adjacency with *DH-ex-Maf/Cck*, whereas excitatory neurons remained spatially distinct from oligodendrocytes and motoneurons—recapitulating the canonical dorsal sensory versus ventral motor organization described in prior molecular and anatomical studies [26].

Myelinating populations (oligodendrocytes, OPCs, astrocytes) formed tight, spatially coherent neighborhoods rather than being dispersed, further indicating accurate recovery of glial domains.

Taken together, these proximity patterns demonstrate that BioLACE reconstructs biologically correct microcircuit relationships and respects laminar boundaries. Importantly, the observed neighborhoods align with known anatomy rather than the homogenized patterns typical of over-smoothing, confirming that the spatial Laplacian and contrastive terms jointly preserve both tissue topology and cell-type specificity.

#### Functional communication sharpened by ligand–receptor topology

To test whether anatomical coherence translates into functional accuracy, we examined 22,745 ligand–receptor (L–R) pairs and observed consistent spatial tightening of biologically established signaling axes: *Bdnf–Ntrk2* (Δ*z*_median_ = −41), *Sema3a–Plxna1* ( −31), and *Sst–Sstr2* ( −20) (Fig. 5d). These shifts recapitulate canonical spinal motifs—neurotrophic support in dorsal sensory zones (*Bdnf–Ntrk2* ), commissural guidance networks (*Sema3a/ 3f–Plxna1/A3* ), and local inhibitory neuromodulation (*Sst–Sstr2* )—and show that BioLACE pulls sender–receiver populations into realistic spatial proximity. The resulting communication map is more compact and biologically interpretable, indicating that BioLACE preserves the functional topology of cell–cell signaling rather than merely enforcing geometric smoothness.

#### Summary

Across the MERFISH spinal cord dataset, integrating curated marker-gene priors enables BioLACE to jointly recover spatial structure and molecular identity. The inclusion of biological priors substantially increases spatial modularity, sharpens boundaries among major neuronal, glial, and vascular populations, and improves alignment with known laminar and perivascular organization. These anatomically coherent embeddings yield more compact and interpretable ligand–receptor topologies, with sender–receiver pairs contracting along established signaling axes. Taken together, the results demonstrate that embedding biological knowledge directly into the learning objective produces spatial representations with verifiable structural and functional validity.

### 3.3 Ablation study on MERFISH hypothalamus reveals the contributions of spatial and biological objectives

#### Overview and significance

Supplementary Fig. 3 summarizes how each objective contributes to clustering quality in the MERFISH hypothalamus. The VAE backbone captures global transcriptomic variation but lacks geometric specificity. Adding Laplacian regularization improves local spatial continuity yet over-smooths adjacent neuronal groups, while contrastive learning alone increases molecular discrimination but fragments spatial gradients. Only their calibrated combination yields embeddings that are simultaneously spatially coherent and biologically separable, peaking at ARI 0.62. Supplementary Method 1 provides full design details and qualitative spatial visualizations.

## 4 Discussion and Conclusion

BioLACE provides a unified framework that combines gene expression, spatial adjacency, and curated marker programs into a single latent space, resolving the tension between spatial smoothness and cell-type interpretability in spatial transcriptomics. Together, our results support a broader goal: cell-type embeddings that integrate spatial context with structured biological knowledge. By embedding marker priors directly into the learning process, BioLACE produces latent spaces with clearer cell-type boundaries and more interpretable biological structure, advancing the analysis of high-resolution spatial omics data.

## Supplementary Methods

### 1. Expanded ablation details: MERFISH hypothalamus

#### Motivation

To isolate the contribution of each objective in BioLACE, we ablated the VAE backbone, Laplacian regularization, and marker-aware contrastive term on the MERFISH mouse hypothalamus dataset. The hypothalamus combines compact neuronal nuclei with diffuse glial regions, providing a stringent test of how spatial and biological constraints interact.

#### Stepwise model development

Supplementary Fig. 3 details the effect of each added regularization term in BioLACE, and Supplementary Table 3 details the quantitative analysis and insights. We began with a *vanilla VAE* (ARI 0.42), which captures global transcriptomic structure but lacks geometric awareness, missing small spatially restricted groups. Adding a *contrastive loss* alone maintained comparable performance (ARI 0.41) with slightly sharper boundaries but unstable partitions—biological similarity alone proved insufficient without spatial context. Introducing the *graph Laplacian* (VAE + Laplacian) improved ARI to 0.45 by promoting local smoothness and alignment with tissue neighborhoods but over-smoothed boundaries between adjacent yet distinct types. Combining both spatial and contrastive objectives required temperature tuning: low *τ* =0.1 over-segmented (ARI 0.40), high *τ* =1.0 over-smoothed (ARI 0.47). At *τ* =0.5, the full model achieved ARI 0.62, cleanly recovering compact neuronal domains.

#### Interpretation

The VAE provides a denoised latent manifold; the Laplacian preserves local topology; and the marker-aware contrastive term sharpens biologically meaningful axes. Calibrated jointly, these components resolve the smoothness–separability trade-off: spatial alignment alone merges types, contrastive separation alone fragments gradients, but the combined objective yields anatomically faithful, biologically interpretable clusters.

### 2. Evaluation Metrics for Cell Clustering and Spatial Analysis

#### Clustering concordance metrics

##### Adjusted Rand Index (ARI)

Measures agreement between predicted clusters and reference labels by counting pairwise co-assignments, with a correction for chance. Given two partitions of *N* samples—predicted clusters *{C*_*k*_*}* and reference labels *{L*_*ℓ*_*}*—with contingency table entries *n*_*kℓ*_ = |*C*_*k*_ ∩ *L*_*ℓ*_|, ARI is:

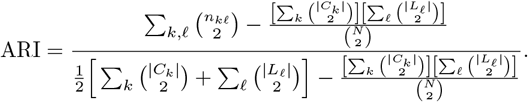

Values range from 0 (chance) to 1 (perfect), with negative values indicating worse-than-random alignment.

##### V-measure

Evaluates cluster quality as the harmonic mean of homogeneity (each predicted cluster contains primarily one true label) and completeness (each true label maps to a single predicted cluster). Let *H*(*·*) denote entropy:

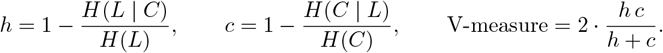

#### Spatial autocorrelation metric

##### Moran’s *I*

Quantifies how strongly gene expression values co-vary among spatial neighbors, detecting whether high or low expression forms spatially contiguous domains. Given expression **g** and spatial weight matrix *W* :

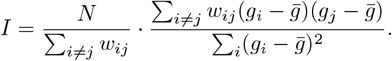

Values near 1 indicate strong spatial clustering.

#### Gene–gene coherence

##### Spearman’s *ρ*

Measures monotonic co-expression between two genes by comparing rank-order agreement:

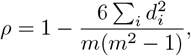

where *d*_*i*_ is the difference in ranks.

#### Binary classification performance

##### Accuracy

Fraction of correctly labeled samples:

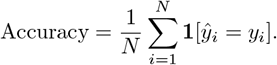

##### Recall

Fraction of true positive samples recovered:

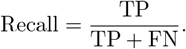

##### AUROC

Probability that a randomly chosen positive example is assigned a higher score than a negative:

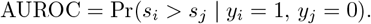

#### Segmentation and spatial-overlap metric

##### Dice Coefficient

Measures spatial overlap between predicted and reference regions: Higher values indicate tighter anatomical correspondence.

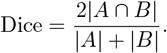

## Supplementary Tables

**Supplementary Table 1.** Slide-seq summary: spatial and functional validation under BioLACE. Metrics use the best BioLACE cluster for each domain.

| Dataset | Domain (cluster) | Markers | Moran's $I$ | AUROC | Within $\rho$ | Dice |
| --- | --- | --- | --- | --- | --- | --- |
| V1 | Purkinje (4) | *Pcp4*, *Pcp2* | 0.263 | 0.948 | 0.649 | 0.699 |
| V1 | Myelin (1) | *Plp1*, *Mbp* | 0.289 | 0.839 | 0.200 | 0.678 |
| V2 | Purkinje (8/2) | *Pcp2*, *Pcp4* | 0.302 | 0.812 | 0.709 | 0.506 |
| V2 | Myelin (5) | *Plp1*, *Mbp* | 0.319 | 0.968 | 0.469 | 0.728 |

**Supplementary Table 2.**
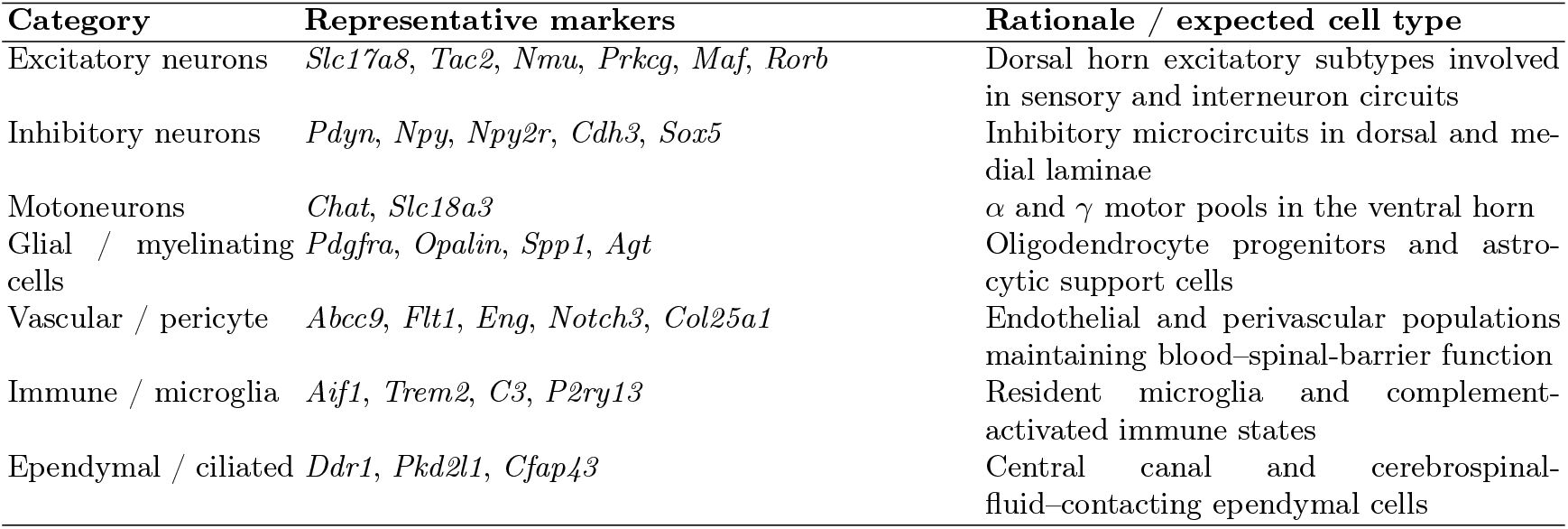
Curated marker genes and rationale for inclusion. Each group represents a biologically distinct program or anatomical niche in the spinal cord.

| Category | Representative markers | Rationale / expected cell type |
| --- | --- | --- |
| Excitatory neurons | *Slc17a8*, *Tac2*, *Nmu*, *Prkcg*, *Maf*, *Rorb* | Dorsal horn excitatory subtypes involved in sensory and interneuron circuits |
| Inhibitory neurons | *Pdyn*, *Npy*, *Npy2r*, *Cdh3*, *Sox5* | Inhibitory microcircuits in dorsal and medial laminae |
| Motoneurons | *Chat*, *Slc18a3* | $\alpha$ and $\gamma$ motor pools in the ventral horn |
| Glial / myelinating cells | *Pdgfra*, *Opalin*, *Spp1*, *Agt* | Oligodendrocyte progenitors and astrocytic support cells |
| Vascular / pericyte | *Abcc9*, *Flt1*, *Eng*, *Notch3*, *Col25a1* | Endothelial and perivascular populations maintaining blood–spinal-barrier function |
| Immune / microglia | *Aif1*, *Trem2*, *C3*, *P2ry13* | Resident microglia and complement-activated immune states |
| Ependymal / ciliated | *Ddr1*, *Pkd2l1*, *Cfap43* | Central canal and cerebrospinal-fluid–contacting ependymal cells |

**Supplementary Table 3.**
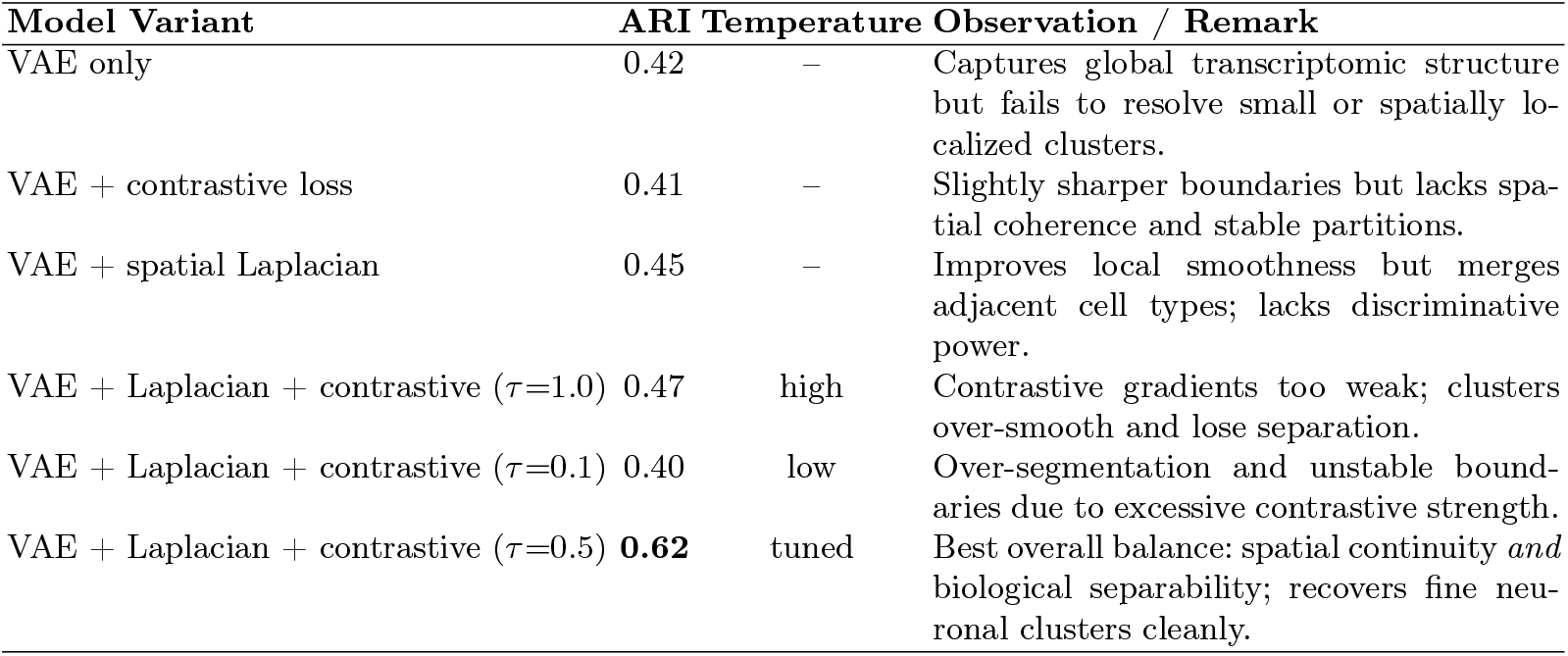
Ablation study on MERFISH hypothalamus. Each component contributes uniquely to clustering performance; only the joint model achieves both spatial and biological coherence.

## Supplementary Figures

**Supplementary Fig. 1.**
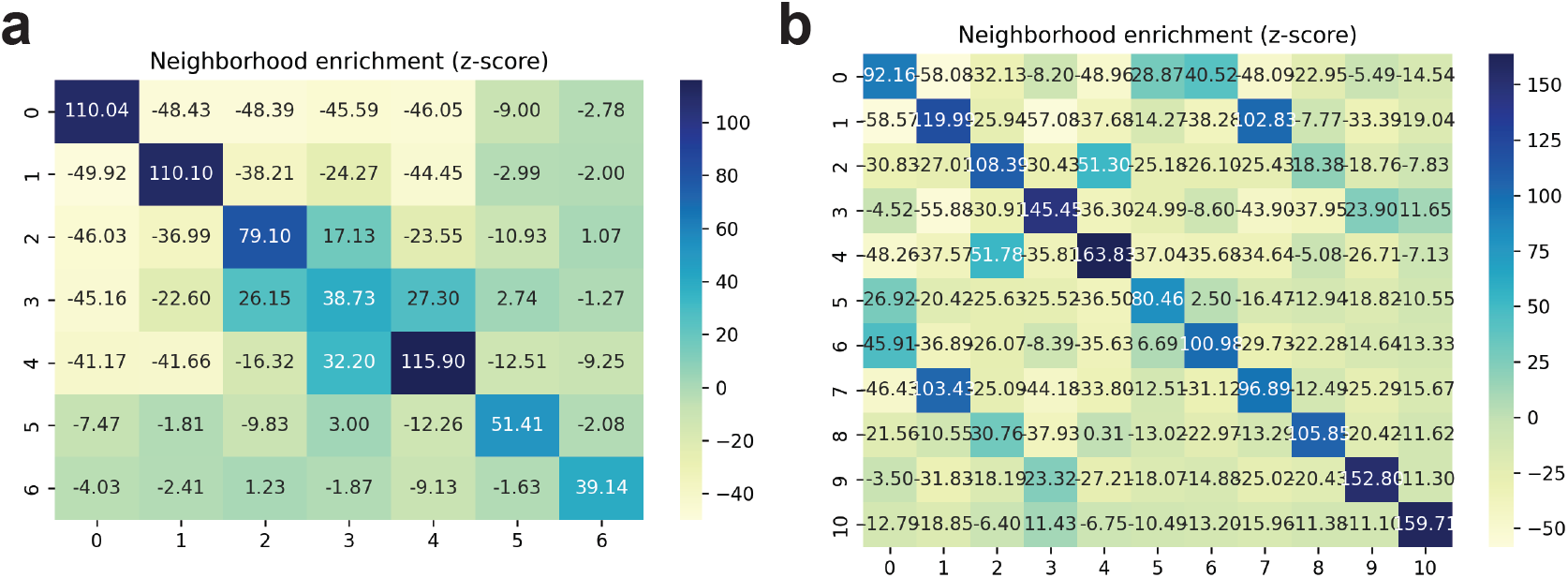
Neighborhood coherence under BioLACE (Slide-seq V1/V2). Squidpy neighbor-enrichment matrices for **(a)** V1 and **(b)** V2. BioLACE shows consistently higher same-label probability among spatial neighbors compared with baselines, indicating that Laplacian alignment preserves local tissue topology without oversmoothing across known boundaries.

**Supplementary Fig. 2.**
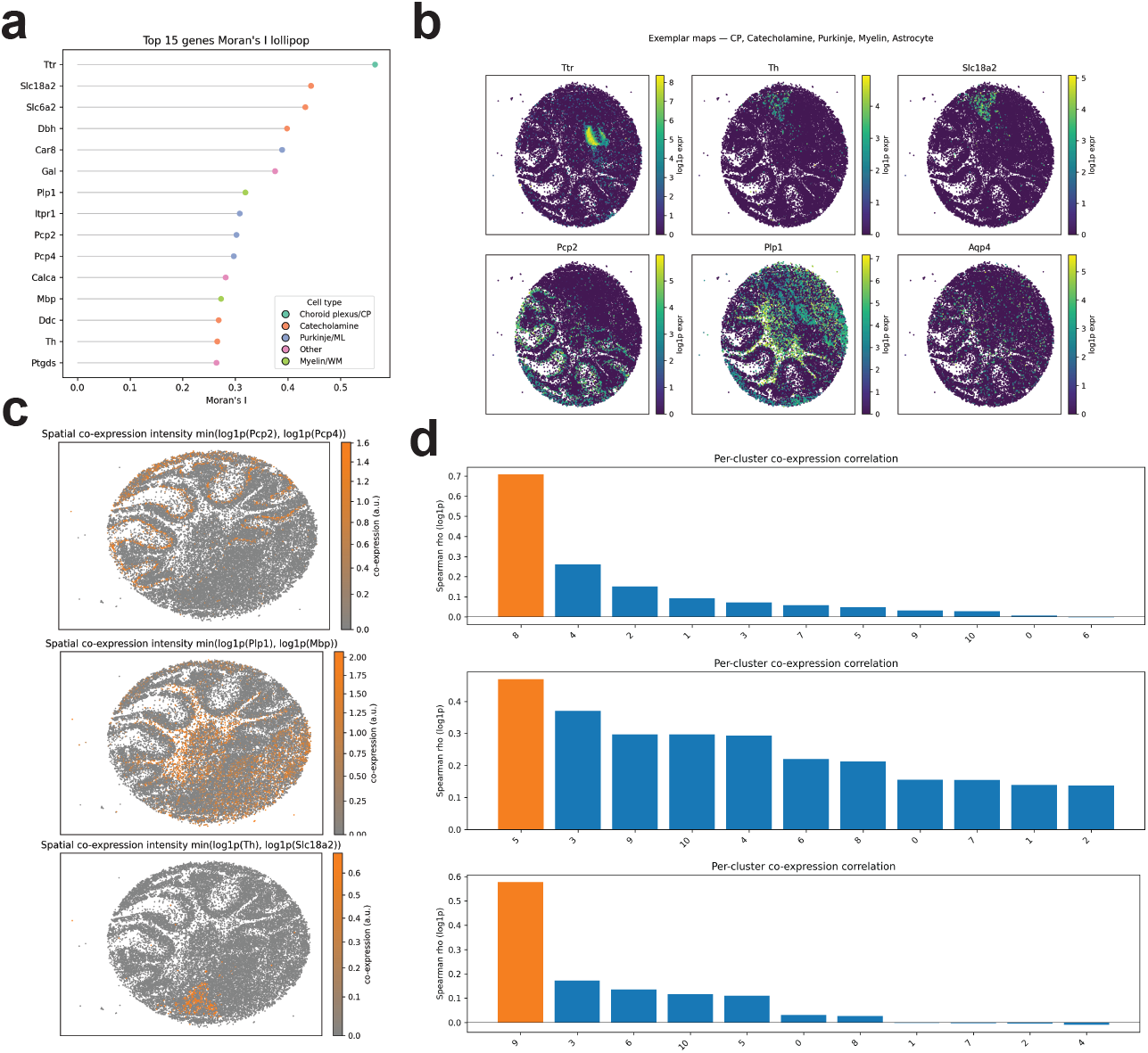
Slide-seq V2—Moran’s *I*, co-expression, and cross-method comparison. **(a)** Lollipop plot of Moran’s *I* identifies top spatially autocorrelated markers: *Plp1* (myelin, *I*=0.319), *Pcp2* (Purkinje, *I*=0.302), and *Ttr* (ependymal/CSF, *I*=0.565). **(b)** Spatial overlays show distinct localization of myelin and Purkinje domains, recapitulating cerebellar architecture. **(c)** Within-cluster co-expression peaks coincide with Moran’s *I* maxima: *Pcp2* –*Pcp4 ρ*=0.709 (Purkinje) and *Plp1* –*Mbp ρ*=0.469 (myelin). **(d)** BioLACE reveals a cluster coexpressing *Th* and *Slc18a2* that is not recovered by RCTD, illustrating added biological resolution.

**Supplementary Fig. 3.**
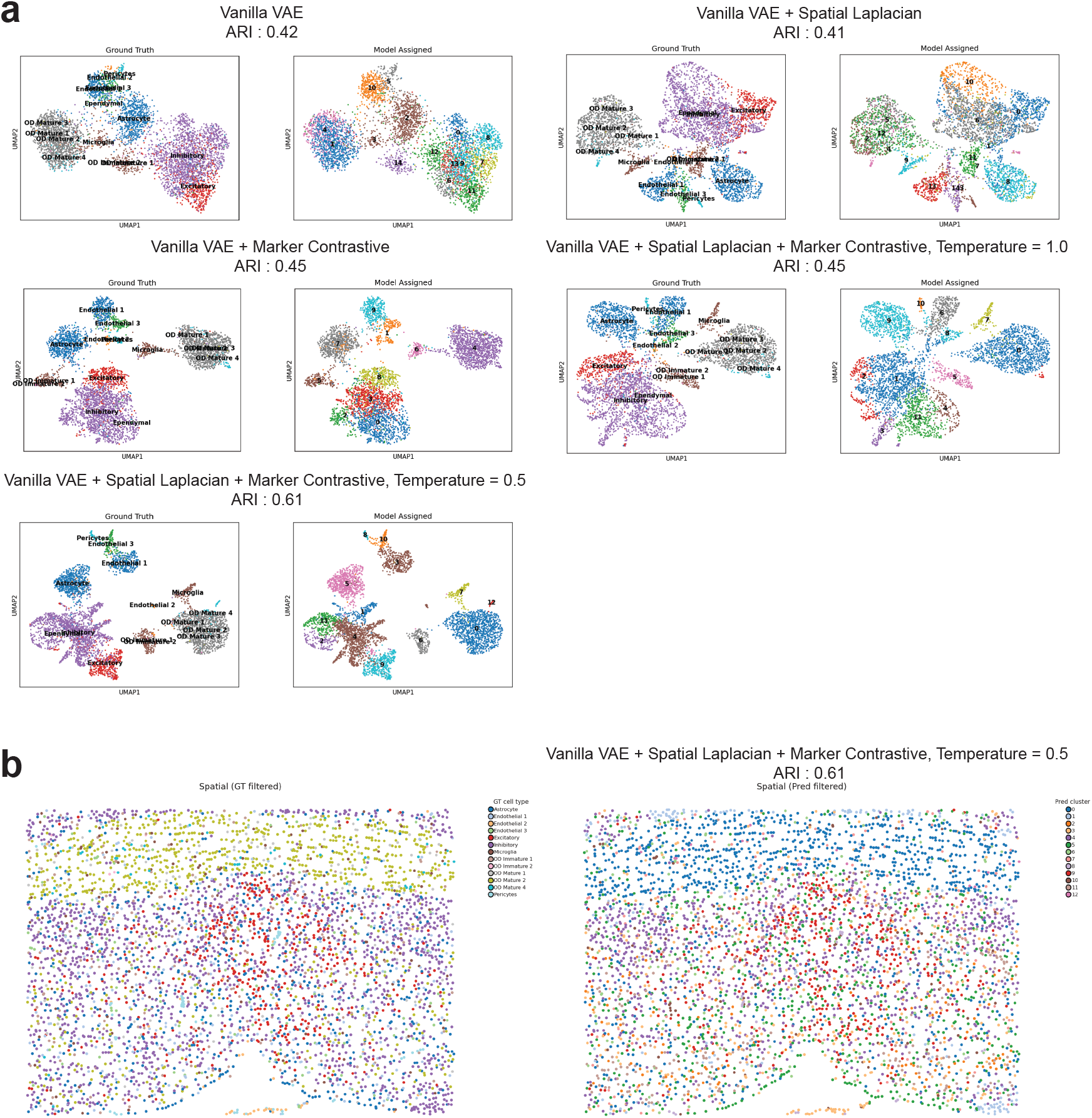
Ablation on MERFISH hypothalamus. **(a)** Progressive inclusion of spatial and biological objectives yields increasingly coherent, anatomically consistent clustering. A vanilla VAE (ARI 0.42) preserves global structure but misses compact nuclei. Adding only contrastive learning (ARI 0.41) sharpens some boundaries but lacks spatial coherence, whereas adding only the Laplacian (ARI 0.45) enforces continuity but merges adjacent types. Joint training requires temperature tuning: high temperature (*τ* =1.0, ARI 0.47) over-smooths; low temperature (*τ* =0.1, ARI 0.40) over-segments. The tuned full model (*τ* =0.5) achieves the best balance (ARI 0.62), recovering compact, anatomically faithful neuronal domains. **(b)** Spatial embeddings (filtered for clusters with ≥ 5 cells to exclude small ependymal and mature oligodendrocyte clusters) show that only the full model resolves compact neuronal domains without merging glial boundaries.

